# Developmental onset of a cerebellar-dependent forward model of movement in motor thalamus

**DOI:** 10.1101/2021.06.25.449956

**Authors:** James C. Dooley, Greta Sokoloff, Mark S. Blumberg

## Abstract

To execute complex behavior with temporal precision, adult animals use internal models to predict the sensory consequences of self-generated movement. Here, taking advantage of the unique kinematic features of twitches—the brief, discrete movements of active sleep—we captured the developmental onset of a cerebellar-dependent internal model. Using rats at postnatal days (P) 12, P16, and P20, we compared neural activity in two thalamic structures: the ventral posterior (VP) and ventral lateral (VL) nuclei, both of which receive somatosensory input but only the latter of which receives cerebellar input. At all ages, twitch-related activity in VP lagged behind movement, consistent with sensory processing; similar activity was observed in VL through P16. At P20, however, VL activity precisely mimicked the twitch itself, a pattern of activity that depended on cerebellar input. Altogether, these findings implicate twitches in the development and refinement of internal models of movement.

## Introduction

Functional behavior entails the production of complex and precisely timed movements. Proper execution of these movements requires that motor commands are coordinated with movement-related sensory feedback (or reafference; Brooks and Cullen, 2019; Ebbesen et al., 2018; Sathyamurthy et al., 2020). This coordination is made possible by the production of copies of motor commands (or corollary discharges) that contribute to the computation of a forward model in the cerebellum and other structures, providing the animal with an internal representation of its own movements (Bastian, 2006; Crapse and Sommer, 2008; Franklin and Wolpert, 2011; Hull, 2020). One of the major targets of the cerebellum is the motor thalamus—the ventral lateral nucleus (VL)—which projects to primary motor cortex (M1; Bosch-Bouju et al., 2013; Dacre et al., 2021; Gornati et al., 2018; Habas et al., 2019; Sauerbrei et al., 2020; Wolpert et al., 1998). In its entirety, this system enables flexible and adaptive updating of movements in real time.

To understand how internal representations of movement in the cerebellum contribute to behavior, researchers have assessed either learning-related changes in cerebellar activity as an animal learns a novel behavioral task (Bosch-Bouju et al., 2014; Gaidica et al., 2018; Wagner et al., 2019) or the behavioral consequences of manipulating the cerebellothalamic system (Dacre *et al*., 2021; Nashef et al., 2019; Sauerbrei *et al*., 2020). Although these experimental approaches have effectively revealed cerebellar functioning in adults, they are not suitable for addressing the question of how and when internal representations first develop. Here, we answer this question by exploiting the unique kinematic features of myoclonic twitches, a self-generated movement that is performed abundantly throughout early development during active (or REM) sleep.

In infant rats, twitches exhibit several features that make them well-suited for directing sensorimotor development (Blumberg et al., 2013a; Blumberg et al., 2020). Specifically, twitches are brief and discrete, hundreds of thousands of them are produced each day, and twitches reliably trigger reafferent activity throughout the sensorimotor system. In addition, the motor command that triggers a twitch also produces a corollary discharge that is conveyed to the cerebellum via precerebellar nuclei (Mukherjee et al., 2018). Here, we hypothesize that twitches also contribute to the development of a forward model of movement to enable comparisons between expected and actual feedback. Because the cerebellar system is still undergoing substantial anatomical and functional development between P12 and P20 (Altman, 1969; Freeman and Nicholson, 2000; 2004; Nicholson and Freeman, 2003; Watanabe and Kano, 2011), a heretofore untested premise of our hypothesis is that twitches continue to contribute to sensorimotor development at ages beyond those that we have previously investigated.

Here, using P12, P16, and P20 rats, we examined twitch-related activity in VL and M1, as well as the thalamic ventral posterior nucleus (VP). First, we demonstrate that twitches, from P12 to P20, continue to drive topographically precise neural activity in all three structures. Next, we show that VP and VL exhibit temporal sharpening of twitch-related activity such that, by P20, the duration of the neural activity closely matches the duration of the twitch. Moreover, whereas twitch-related activity in VP reliably lags movement at all three ages—as expected of a sensory nucleus—we find no such lag in VL activity at P20. In fact, at P20, VL activity precisely mimics the twitch itself. Given that VL receives strong, direct projections from the deep cerebellar nuclei (DCN), we pharmacologically inactivated the DCN at P20 and demonstrate that the precise co-occurrence of VL activity with movement is disrupted. Altogether, these results demonstrate that, by P20, twitch-related activity in VL reflects the emergence of a cerebellar-dependent capacity to accurately predict the sensory consequences of movement.

## Results

### Sleep-related twitches continue to drive thalamocortical activity

To establish that twitches continue to drive thalamocortical activity beyond early infancy, we recorded neural activity in head-fixed, unanesthetized rats at P12, P16, and P20. Rats were tested in the Mobile HomeCage, which allows animals to locomote while head-fixed (Fig. 1A, left). Extracellular activity was recorded in the forelimb region of M1 (P12: N = 11 pups, 121 neurons; P16: N = 12 pups, 177 neurons; P20: N = 22 pups, 259 neurons), VP (P12: N = 6 pups, 116 neurons; P16: N = 6 pups, 82 neurons; P20: N = 5 pups, 75 neurons) and VL (P12: N = 5 pups, 57 neurons; P16: N = 5 pups, 89 neurons; P20: N = 6 pups, 89 neurons; Fig. 1A, right; Fig. S1A). Neural activity, electromyographic (EMG) activity in the nuchal and biceps muscles, and high-speed video (100 frames/s) were recorded continuously for 3-6 hours. Representative behavioral and neurophysiological data during a 15-s period of active sleep at P12, P16, and P20 are presented in Fig. 1C. Rats cycled between sleep and wake at all three ages and, as expected, the percentage of time spent in active sleep decreased across age (Fig. 1B, S1B; see Frank and Heller, 1997; Gramsbergen et al., 1970; Jouvet-Mounier et al., 1970). Along with a significant age-related decrease in total time in active sleep during each recording session (Fig. S1C), older pups also showed significantly fewer twitches (Fig S1D).

**Figure 1:**
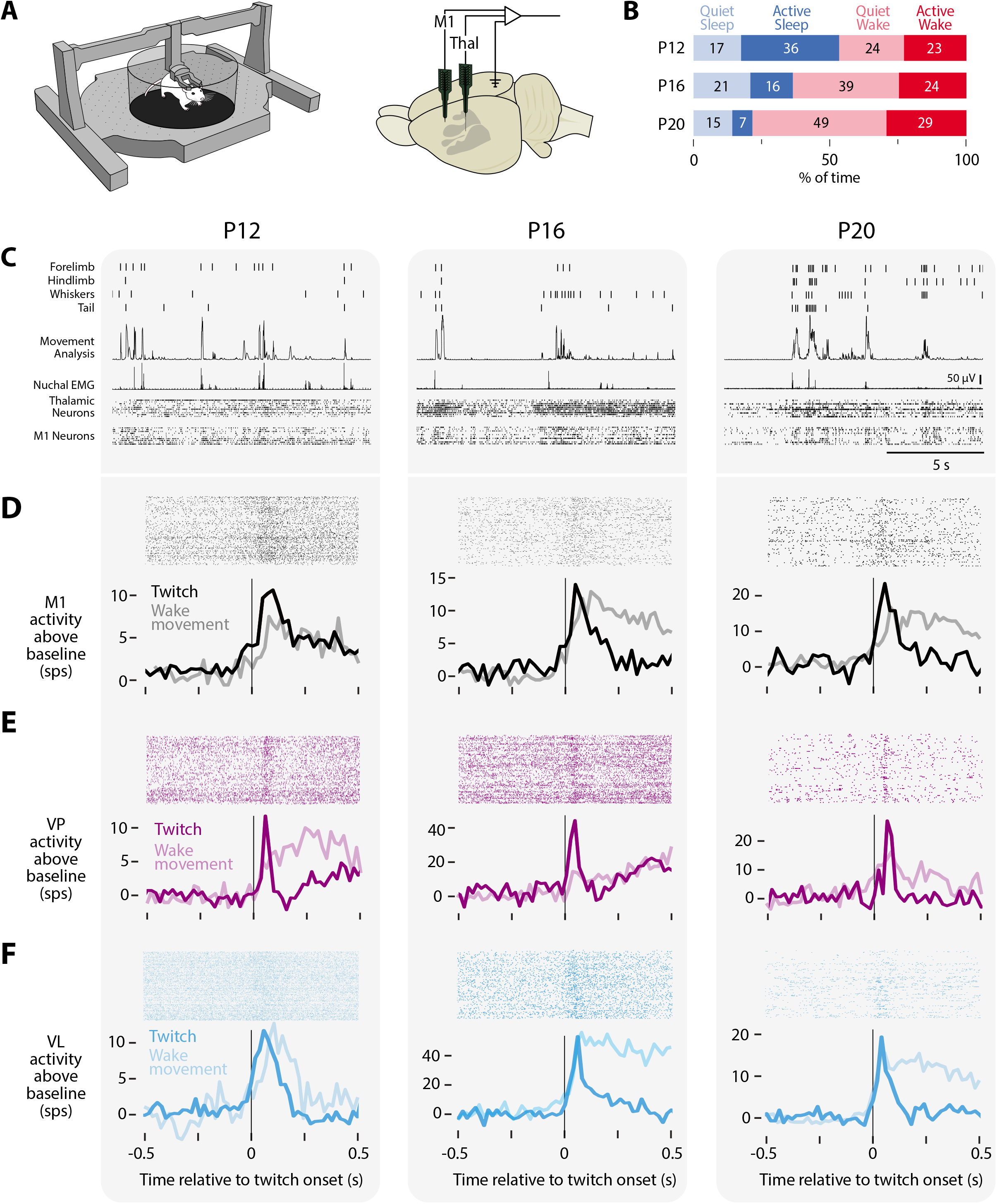
Twitch-related reafference continues to trigger thalamocortical activity. (A) Left: An illustration of the Mobile HomeCage used to collect behavioral and electrophysiological data across sleep and wake from head-fixed rats. Right: An infant rat brain, with primary somatosensory cortex denoted in gray, showing the relative locations of the recording sites in M1 and thalamus, as well as the location of the reference/ground. (B) Mean percentage of time spent in quiet sleep (light blue), active sleep (dark blue), quiet wake (light red), and active wake (dark red) at P12, P16, and P20. Numbers indicate the percentage of time for each category. (C) Representative 15-s behavioral and electrophysiological records at P12, P16, and P20. From top: twitches of four different body parts, the amount of movement detected in each video frame, rectified nuchal EMG, and thalamic and M1 neural activity (each row is a different neuron). (D) Top: Raster sweeps for representative twitch-responsive M1 neurons at P12, P16, and P20, with each row triggered on a different twitch. Bottom: Perievent histograms (bin size = 10 ms) showing the neuron’s firing rate above baseline triggered on twitches (dark lines) and wake movements (light lines). (E) Same as in (D), except for VP. (F) Same as in (D), except for VL. See also Figure S1.

At P12, P16, and P20, neurons in M1 increased their activity only after twitches (Fig. 1D, black), similar to previous findings through P12 and consistent with the view that M1 is not involved in motor control at these ages (Dooley and Blumberg, 2018; Singleton et al., 2021; Young et al., 2012). Twitch-responsive neurons in M1 also responded to wake movements (Fig. 1D, gray). Although such responses are presumed to arise from thalamus, the thalamic source of sensory inputs to M1, particularly in early development, remains unclear (Gómez et al., 2021). As with M1, neurons in VP and VL also showed increased activity in response to twitches and wake-movements at all three ages (Fig. 1E, F). At P12 and P16, twitch-responsive neurons in VP exhibited either a fast response to twitches (with a rapid return to baseline), a slow response (with neural activity elevated for more than 2 s), or a fast-and-slow response comprising two distinct peaks in activity (Fig. S1E). However, by P20, the slow response disappeared, leaving only neurons with the fast response (Fig. S1F). The existence of a slow response in VP is reminiscent of similarly transient long-duration responses to brief light flashes in the visual system, which result from cortical feedback to thalamus (Colonnese et al., 2010; Murata and Colonnese, 2018). In contrast, all VL neurons exhibited a fast response regardless of age (Fig. S1G-H). The absence of slow responses in VL neurons suggests a difference from VP in its development of corticothalamic feedback.

### Somatotopic and temporal refinement of twitch-related thalamic activity

Having demonstrated that neurons in VP and VL exhibit twitch-related activity through P20, we next assessed whether this activity is somatotopically and temporally precise. With respect to somatotopy, we tracked twitching across four body parts (forelimb, hindlimb, whiskers, and tail; Fig. S2A-B) and then characterized neural responses in VP and VL to twitches of each part. Neurons in both structures responded with somatotopic precision across all three ages: When a neuron was responsive to twitches of one body part (e.g., forelimb), it did not respond to twitches of other body parts (e.g., hindlimb, whiskers, or tail; Fig. 2A-B; see also Fig. S2C).

**Figure 2:**
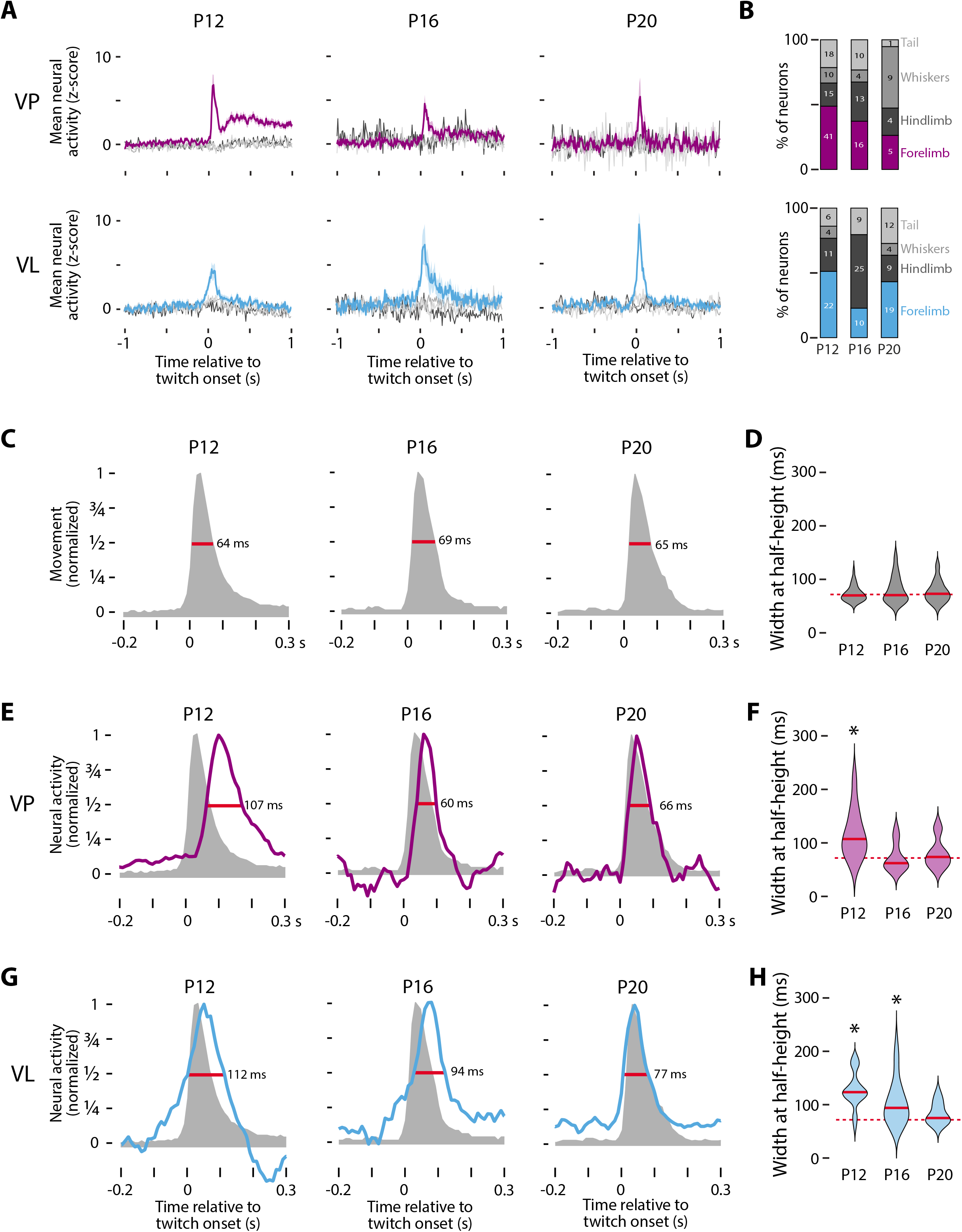
Twitch-related thalamic activity is spatiotemporally refined across age. (A) Mean (±SE) z-scored neural activity of forelimb twitch-responsive neurons in VP (top) and VL (bottom) in response to twitches of the forelimb (purple or cyan, respectively), hindlimb (dark gray), whiskers (gray), or tail (light gray). (B) Percentage of twitch-responsive neurons that respond to twitches of different body parts in VP (top) and VL (bottom). The number of neurons for each category is also shown. (C) Representative normalized movement profiles for forelimb twitches at P12, P16, and P20. The widths at half-height (red lines) are also shown. (D) Violin plots of the width at half-height for twitch displacent at each age. The solid red line is the median value at each age and the dotted red line is the grand median across all ages. (E) Representative twitch-related responses of VP neurons at P12, P16, and P20 (blue). The movement profiles from (C) are shown for comparison. The widths at half-height (red lines) are also shown. (F) Violin plots of the width at half-height for twitch-responsive VP neurons at each age; the solid red lines are the median value at each age. Asterisks indicate significant difference (P < 0.0167) between the median value for the neural response and the grand median duration of the twitch (dotted red line). (G) Same as in (E), except for VL. (H) Same as in (F), except for VL. See also Figure S2.

To assess the temporal precision of thalamic activity, we first characterized the kinematics of twitches at each age. Twitches are rapid movements, with maximum durations of <100 ms (or fewer than 10 frames; Fig. S2A). Generally, twitches are characterized by rapid acceleration to a peak displacement, followed by a slow return to baseline (Fig. 2C). This general shape was consistent across body parts and ages, with peak displacement occurring 30 ms (or three frames) after the twitch was triggered (Fig. S2D). In contrast, wake movements exhibited more variability in peak displacement and were over ten times longer in duration (Fig. S2D). To estimate the duration of each twitch, we determined the width at half-height of the displacement curves for all body parts. There was no significant age-related change in mean twitch duration (H_(2,143)_ = 0.82, P = 0.66). These results are not specific to half-height, as we found similar results using the width at both one-quarter and three-quarter height (data not shown).

We also used width at half-height to compare the duration of the neural responses to twitches in VP and VL. To ensure we were making a comparison across similar populations of neurons, we restricted this analysis to the “fast” VP neurons (see Fig. S1E). In VP, there was a significant effect of age on response duration (H_(2,55)_ = 16.3, P < 0.001), with durations at P12 being longer than those at P16 and P20 (Fig. 2E, F). Furthermore, by comparing the duration of the VP response with the duration of twitches, we found a significant difference at P12 (Z_28_ = 4.43, P < 0.001), but not at P16 or P20 (P16: Z_18_ = - 0.57, P = 0.57; P20: Z_9_ = 0.31, P = 0.79). Thus, by P16, the increase in neural activity in VP accurately represents the temporal duration of twitches.

Parallel analyses in VL also revealed a significant effect of age on response duration, with durations decreasing with age (Fig. 2G; H_(2,63)_ = 18.2, P < 0.001). Unlike VP, in VL we saw a continuous decrease in response duration at all three ages, with neural responses at P12 and P16, but not P20, exhibiting durations that were significantly longer than those of twitches (Fig. 2H; P12: Z_24_ = 4.20, P < 0.001; P16: Z_23_ = 3.36, P < 0.001; P20: Z_22_ = 1.81, P = 0.07). Again, these results in VP and VL were not specific to width at half-height, as the same basic results held at both one-quarter and three-quarter height (data not shown).

### Divergent representations of movement in VP and VL

As demonstrated thus far, by P20, twitch-related neural activity in VP and VL is somatotopically precise and matches the duration of twitches. Next, we examined the difference in the timing of the peak neural responses in relation to peak twitch displacement. We then classified movement-related neural activity as follows: activity that reliably preceded movement was classified as “motor” (Fig. 3A, red); neural activity that reliably followed movement was classified as “sensory” (Fig. 3A, green); and neural activity that reliably co-occurred with movement was classified as “internal model” (Fig. 3A, blue; see Gaidica *et al*., 2018).

**Figure 3:**
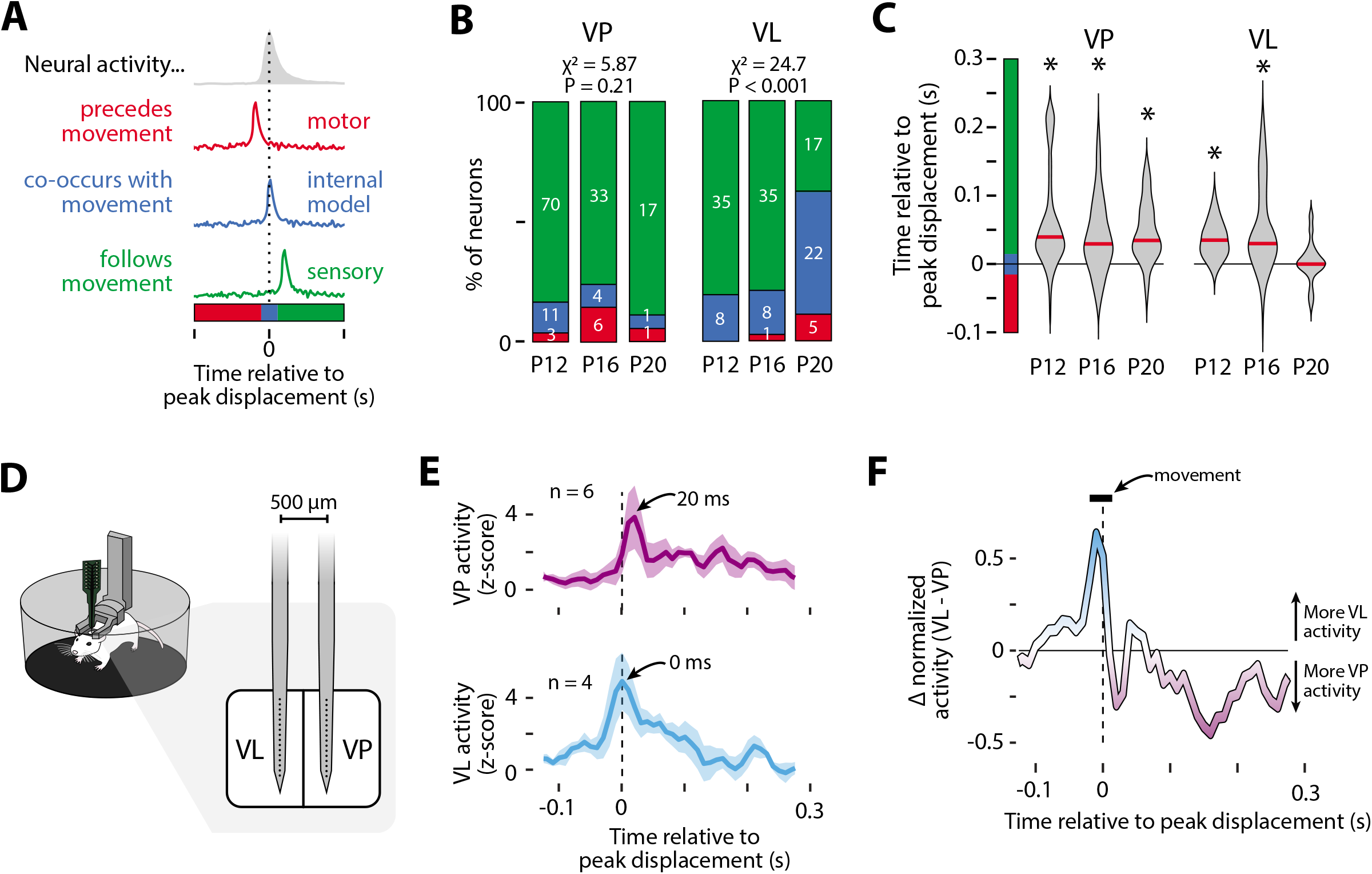
Diverging representations of movement in VP and VL. (A) Illustration of the possible temporal relationships between twitches and neural activity: motor (red), internal model (blue), and sensory (green). (B) Percentage of neurons belonging to each category in (A). The number of neurons for each category is also shown. The percentage of neurons in each category was significantly different across age for VL (P < 0.001) but not VP. (C) Violin plots of the time relative to peak displacement for VP (left) and VL (right) neurons at each age. Asterisks indicate that the median value (red line) is significantly different (P < 0.0167) from 0. (D) Illustration of the electrode used to perform dual recordings in VL and VP in P20 rats. (E) Mean (±SE) z-scored neural activity of forelimb twitch-responsive neurons in VP (top) and VL (bottom) relative to peak forelimb displacement (dotted lines). Data are from one representative pup. The number of neurons contributing to each plot are shown. (F) Difference between VL and VP activity as calculated from the normalized perievent histograms in (E), showing that VL exhibits more twitch-related activity than VP before peak displacement, whereas VP shows more twitch-related activity than VL after peak displacement (dotted line). The duration of the median movement (horizontal bar) is also shown. See also Figure S3.

VP’s role in transmitting somatosensory information to cortex is well-established (Ebner and Kaas, 2015). Consequently, we expected twitch-related activity in VP to reflect sensory feedback from twitches, and this was exactly what we saw. We categorized over 80% of VP neurons as sensory at all three ages, with no significant change in group identity across age (Fig. 3B; χ^2^ = 5.87, N = 146, P = 0.21; see also Fig. S3A). Further, at all ages, twitch-related activity in VP significantly lagged peak twitch displacement by 30-40 ms (Fig. 3C; P12: Z_73_ = 6.69, P < 0.001; P16: Z_38_ = 4.09, P < 0.001; P20: Z_16_ = 3.16, P < 0.01), consistent with the conduction delay of twitch reafference found in other somatosensory structures (Dooley and Blumberg, 2018).

In adults, the available evidence suggests that VL is not strictly sensory like VP. For example, VL neurons can selectively respond to movements that are self- or other-generated (Butler et al., 1992a; Butler et al., 1992b), with the activity of neurons responsive to self-generated movements being too fast to be sensory (Bosch-Bouju *et al*., 2014; Gaidica *et al*., 2018). Other studies suggest that VL can influence the production of movement via its projections to M1 (Dacre *et al*., 2021; Sauerbrei *et al*., 2020). Here, at P12 and P16, most VL neurons were classified as sensory (Fig. 3B) because of a significant 30-40-ms lag in peak neural activity in relation to peak displacement (Fig. 3C; P12: Z_38_ = 6.69, P < 0.001; P16: Z_39_ = 4.09, P < 0.001). By P20, however, most VL neurons did not exhibit a lag (Fig. 3B, right; χ^2^ = 24.7, N = 131, P < 0.001; Fig. 3C, right; Z_35_ = 0.676, P = 0.500).

Until now, we recorded VP and VL activity separately in different animals. Thus, to increase our confidence in the differential timing of twitch-related activity in VP and VL, we performed simultaneous recordings in the two nuclei in an additional set of P20 rats (n = 3; Fig. 3D). Focusing on neurons that responded to forelimb twitches, we again found that VP activity lagged peak limb displacement by 20 ms, with VL activity exhibiting no lag at all (Fig. 3E-F). Given that the sensory input to VP arrives directly from the dorsal column nuclei (Lund and Webster, 1967; Makous et al., 1996), the fact that twitch-related activity occurred in VL 20 ms before that in VP indicates that the VL activity is unlikely to be due to reafference.

### Disrupting cerebellar output at P20 unmasks sensory responses in VL

If twitch-related VL activity at P20 reflects the emergence of a forward model, we would expect the cerebellum to be the source of this activity for several reasons. First, the deep cerebellar nuclei (DCN) are the dominant source of excitatory input to VL (Habas *et al*., 2019); and second, the cerebellum is responsible for the precise timing of movement-related activity in VL (Gaidica *et al*., 2018; Nashef *et al*., 2019). Accordingly, we predicted that inactivation of the DCN at P20 would disrupt the precise timing of VL activity, resulting in activity that resembles sensory responses in VP. To test this prediction, we recorded activity in VL after injecting 0.5 μL of muscimol (N = 6 pups; 96 VL neurons) or saline (N = 6 pups; 93 VL neurons) into the DCN (Fig. 4A). Injections targeted the interposed nucleus, which conveys input to VL (Fig. S4A; Quy et al., 2011). DCN inhibition had no effect on the percentage of time spent in active sleep (Fig. S4B), though it did cause a small but significant decrease in VL activity (Fig. S4C; F_(1,152)_ = 8.42, P < 0.005). DCN inhibition also had no effect on the rate of twitching (Fig. S4D), the somatotopic distribution of twitch-responsive neurons (Fig. S4E), or the kinematics of twitches (Fig. S4F). There were also no significant group differences in the duration (width at half-height) of twitches or VL activity (Fig. S4G).

**Figure 4:**
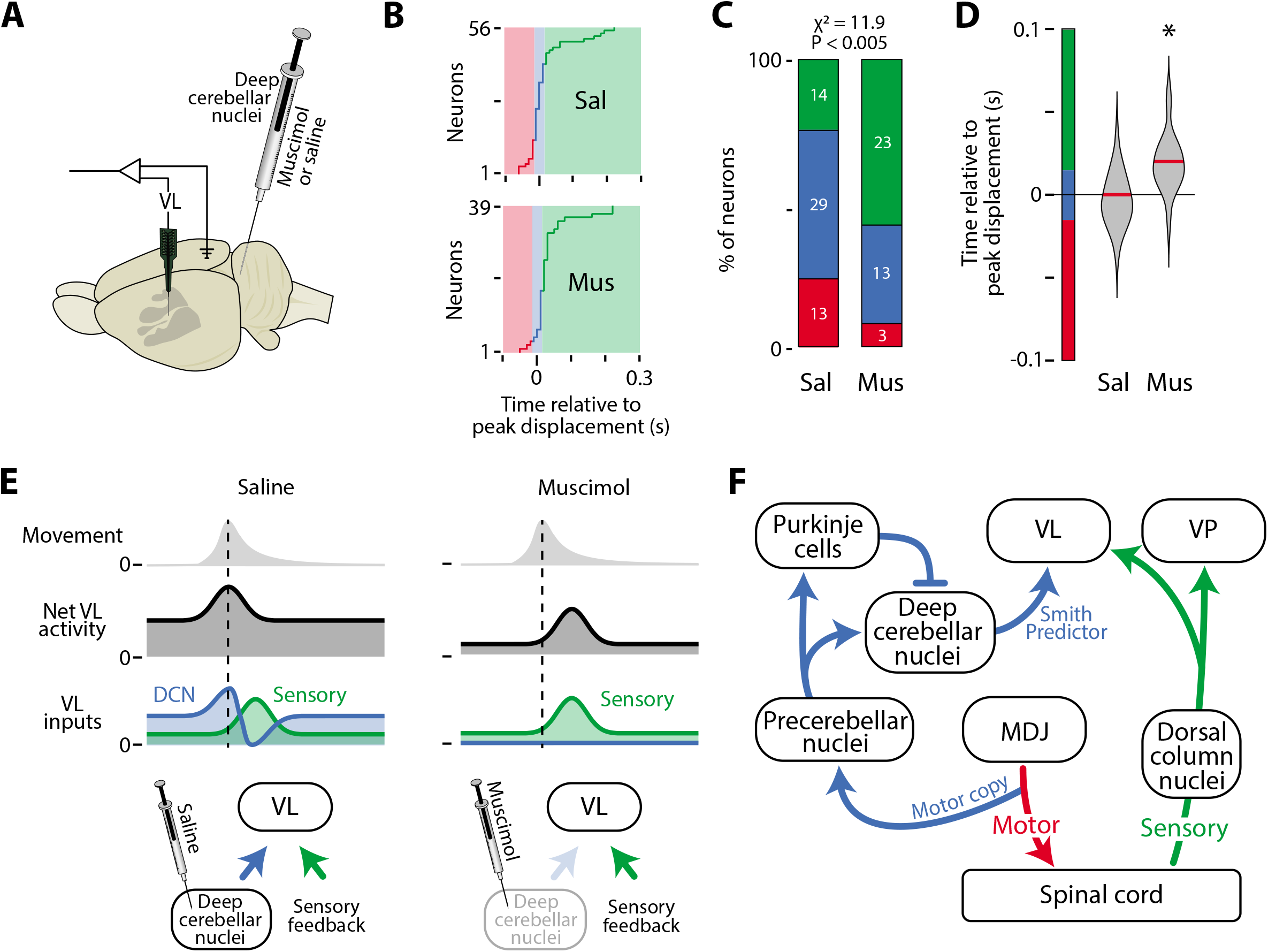
DCN inhibition at P20 disrupts VL activity. (A) Illustration of an infant rat brain to show the electrode locations and the microsyringe location in the DCN. (B) Timing of peak neural activity relative to peak displacement for all twitch-responsive neurons in the saline (Sal; top) and muscimol (Mus; bottom) groups. Color codes as in Fig. 3A-B. (C) Percentage of neurons in (B) belonging to each category in the Sal and Mus groups. The number of neurons for each category is also shown. The percentage of neurons in each category was significantly different between the two groups (P < 0.005). (D) Violin plots of the time relative to peak displacement of VL neurons in the Sal and Mus groups. Asterisks indicate that the median value (red line) is significantly different from 0 (P < 0.025). (E) Proposed model explaining twitch-related VL activity in the saline (left) and muscimol (right) groups. In saline-injected pups, VL receives input from the DCN (blue) and sensory feedback (green). Early activity from the DCN (blue peak) provides VL with a forward model of the twitch, and the later decrease in activity (blue trough) cancels the sensory signal (green peak). In muscimol-injected pups, both the early- and the late-arriving signals from the DCN are blocked, thereby revealing the sensory feedback signal. (F) Circuit diagram to support the functional hypothesis in (E). Separate pathways are shown for motor signals (red), sensory signals (green), and signals that convey a forward model to VL (blue). MDJ: mesodiencephalic junction. See also Figure S4.

As predicted, DCN inactivation with muscimol shifted the peak of twitch-related activity in VL (Fig. 4B; Fig. S4H-I), resulting in activity in VL that resembled the sensory responses in VP (see Fig. S3A). Whereas most neurons in the saline group showed peak activity consistent with the action of a forward model, the majority of neurons in the muscimol group showed peak activity consistent with sensory processing, a statistically significant shift (Fig. 4C; χ^2^ = 11.9, N = 95, P < 0.005). Altogether, whereas twitch-related activity in VL neurons of muscimol-injected pups exhibited a significant 20-ms lag in peak twitch displacement (Fig. 4D; Z_39_ = 4.04, P < 0.001), no such lag occurred in the saline-injected pups (Z_56_ = −0.11, P = 0.91).

## Discussion

We have demonstrated here that twitch-related activity continues to trigger somatotopically precise neural activity in VP, VL, and M1 beyond the early postnatal period. We have also shown that twitch-related activity in VP and VL is refined during postnatal development such that, by P20, the response duration is as brief as the twitch itself. Moreover, whereas the neural response in VP reliably lags twitches by 20-30 ms, indicative of reafference, the response in VL undergoes a developmental shift—from a reafferent response at P12 and P16 to a response at P20 that closely mimics the time-course of a twitch. This last finding reveals the emergence in VL of a forward model of movement that is cerebellar-dependent, as inactivating the DCN disrupted this precise correspondence between VL activity and twitches. Further, DCN inactivation unmasked a reafferent response that resembled twitch-related activity in VP. Given that forward models are critical for distinguishing self from other, our findings identify the developmental origin of the self-other distinction.

### Why twitches?

Neural representations of movement are inherently noisy (Franklin and Wolpert, 2011). Thus, to extract a signal that accurately characterizes how movements are represented in the brain, experimenters have typically recorded neural activity while an animal repeats a movement dozens or hundreds of times (Bosch-Bouju *et al*., 2014; Gaidica *et al*., 2018; Sauerbrei *et al*., 2020). Getting animals to repeat the same movement requires weeks of training, precluding investigation into when and how these representations develop. To circumvent this problem, we relied here on twitches, hundreds of which are produced spontaneously and discretely in a single recording session. Moreover, twitches exhibit remarkably consistent kinematics. Finally, because twitches are produced during active sleep, they have a high signal-to-noise ratio (Blumberg et al., 2013b). These features of twitching enable direct comparison of movement-related activity across age, providing a window into how neural representations of movement in sensorimotor structures emerge.

We found here that twitch-related activity in VL co-occurs with movement, peaking 20-30 ms before activity in VP. Similarly timed activity has been found in VL across mammalian species. In primates, VL neurons respond faster than VP neurons to self-generated—but not other-generated—movements (Butler *et al*., 1992a; Butler *et al*., 1992b). In rodents, VL shows rapid movement-related activity, although the timing of this activity has not been directly compared to sensory responses in VP (Bosch-Bouju *et al*., 2014; Dacre *et al*., 2021; Gaidica *et al*., 2018; Sauerbrei *et al*., 2020).

We also found here that cerebellar inactivation disrupts the timing of VL activity, paralleling similar findings in adult animals (Nashef *et al*., 2019; Nashef et al., 2018). However, because previous studies have examined cerebellar activity as animals are producing complicated continuous movements (e.g. reaching for food), directly attributing neural activity in VL to any one facet of the movement has been challenging. Further complicating this issue, cerebellar inactivation can change the kinematics of the movements themselves (Becker and Person, 2019; Dacre *et al*., 2021; Nashef *et al*., 2019). Thus, although the cerebellum clearly shapes movement-related activity in VL, fully characterizing the cerebellum’s contribution to this activity has remained elusive. Importantly, we found here that cerebellar inactivation did not change the kinematics of twitches (see Fig. S4F, G), thereby allowing us to directly compare twitch-related activity in VL with and without cerebellar input. Upon inhibition of DCN activity, we observed a significant delay in twitch-related activity that, we surmise, was caused by the blockade of biphasic DCN input to VL (Fig. 4E, left). Consequently, by blocking DCN output, we unmasked VL activity that arises from the sensory periphery (Fig. 4E, right).

### Is the P20 cerebellum a Smith Predictor?

Applying the principles of mechanical engineering to neuroscience, Miall and colleagues (Miall et al., 1993) hypothesized that the cerebellum functions as a predictive controller called a Smith Predictor. A Smith Predictor comprises two internal models: (1) a simple forward model that mimics the timing of a self-generated movement, enabling prediction of the expected sensory consequences of movement, and (2) a lagged inhibitory copy of the original forward model that accounts for feedback delays in the system (e.g., conduction delays from muscle to brain). When the actual and predicted sensory feedback signals match, they cancel each other out, resulting in no change in overall activity (Fig. 4E, left). We propose that, by P20, the biphasic cerebellar input to VL functions as a Smith Predictor (see Fig. 4F). Accordingly, without cerebellar input to VL, its activity (like that of VP) would predominantly reflect sensory input, as found here.

Previous research in rats supports the above interpretation. In adult rats, DCN neurons show a biphasic response to whisker movements, composed of an early burst of activity at movement onset followed by a longer decrease in activity that results from DCN inhibition by Purkinje cells (Brown and Raman, 2018). Further, in P8 rats, copies of motor commands, including those that produce twitches, are conveyed from the red nucleus and adjacent neurons to precerebellar nuclei (i.e., the inferior olive and lateral reticular nucleus), before projecting to the cerebellum via climbing and mossy fibers (Mukherjee *et al*., 2018). Also, as demonstrated in P8 and P12 rats, the both Purkinje cells and the DCN exhibit clear twitch-related activity (Del Rio-Bermudez et al., 2016; Sokoloff et al., 2015a; Sokoloff et al., 2015b). Thus, we propose that a biphasic response in VL results from two temporally segregated signals from the DCN: an early response that reflects forward-model computations built upon motor copies arising from precerebellar nuclei, and a late response that reflects the lagged inhibitory copy that arises from inhibition of the DCN by Purkinje cells (Fig. 4F).

### Twitches are ideally suited to calibrate a Smith Predictor

The establishment and maintenance of a functioning Smith Predictor relies heavily on accurate timing: To properly cancel the actual sensory feedback from a movement, the delayed inhibition of reafference must precisely match the conduction delay of the reafferent signal. Further complicating this issue, changes in myelination and axonal length during bodily growth and development will modify the conduction delays of sensory feedback (Downes and Mullins, 2014). Accordingly, this system cannot be prespecified or genetically programmed. It must develop.

Miall and colleagues (1993) were aware of this problem and proposed a potential solution: “The size of the feedback time delay could be estimated by measuring the delay between issuing a motor command and assessing its result. This would be most easy to do if the motor command were discrete…, for the reafferent signal would then change abruptly” (Miall *et al*., 1993). Twitches are exactly this type of discrete signal. In other words, Miall and colleagues foreshadowed a solution, based on design principles, that is readily available as a naturally occurring behavior in infant animals. Moreover, given the present demonstration that twitches drive neural activity in thalamic and cortical structures far beyond the period of early infancy, it seems likely that twitches can continue to calibrate the sensorimotor system across the lifespan to enable precise predictions of the expected delays of reafference from wake movements. In this way, twitches and wake movements can be viewed as complementary modes of activation for the sensorimotor system.

## Acknowledgments

We thank Ryan Glanz for technical assistance. This research was supported by grants from the National Institutes of Health (R37-HD081168 to M.S.B. and T32-NS101858 to J.C.D.)

## Author Contributions

Conceptualization, J.C.D., G.S., and M.S.B.; Methodology, J.C.D., G.S., and M.S.B.; Software: J.C.D.; Formal Analysis: J.C.D.; Investigation: J.C.D.; Data Curation: J.C.D.; Writing – Original Draft, J.C.D. and M.S.B.; Writing – Review & Editing, J.C.D., G.S., and M.S.B.; Visualization: J.C.D., G.S., and M.S.B.; Funding Acquisition, J.C.D., G.S., and M.S.B.; Resources, G.S. and M.S.B.; Supervision, G.S. and M.S.B.

## Declaration of Interests

The authors declare no competing interests.

## STAR METHODS

### RESOURCE AVAILABILITY

#### Lead contact

Further information and requests for resources should be directed to, and will be fulfilled by, the lead contact, Dr. James Dooley.

#### Materials Availability

This study did not generate new unique reagents.

#### Data and Code Availability

Both scripts and data in this paper are available upon request.

### EXPERIMENTAL MODEL AND SUBJECT DETAILS

For the first experiment, cortical and thalamic activity was recorded from Sprague-Dawley Norway rats (*Rattus norvegicus*) at P12 (N = 11 pups; 31.0 ± 2.1 g; 3 male), P15-16 (hereafter P16; N = 12 pups; 44.0 ± 3.1 g; 8 male), and P19-23 (hereafter P20; N = 22 pups; 61.6 ± 7.6 g; 15 male). Animals within the same age group were always selected from different litters. Recordings were performed in the forelimb region of M1 and either the ventral lateral (VL) or the ventral posterior (VP) nucleus of thalamus. For an animal to be included in the analysis, it needed more than 40 twitches of a single body part and at least one neuron had to be classified as twitch responsive, criteria that lead to the exclusion of 8 animals. An additional 3 rats were used at P20 for dual recordings in VL and VP. For the experiment in which the deep cerebellar nuclei (DCN) were inactivated, a total of 12 male and female P20 rats were used (n = 6 per group).

Pups were born and reared in standard laboratory cages (48 × 20 × 26 cm) in a temperature- and humidity-controlled room on a 12:12 light-dark cycle, with food and water available ad libitum. The day of birth was considered P0 and litters were culled to eight pups by P3. All pups had at least four littermates until P12 and at least two littermates on the day that neurophysiological recordings were performed. No pups were weaned before testing. All experiments were conducted in accordance with the National Institutes of Health (NIH) Guide for the Care and Use of Laboratory Animals (NIH Publication No. 80–23) and were approved by the Institutional Animal Care and Use Committee of the University of Iowa.

### METHOD DETAILS

#### Surgery

As described previously (Dooley and Blumberg, 2018), a pup with a visible milk band (P12) or a healthy body weight (P16 and P20) was removed from the litter and anesthetized with isoflurane gas (3–5%; Phoenix Pharmaceuticals, Burlingame, CA). The hair on the top of the head was shaved and care was taken to ensure that the vibrissae were intact. Two custom-made bipolar hook electrodes (0.002 inch diameter, epoxy coated; California Fine Wire, Grover Beach, CA) were inserted into the nuchal and biceps muscles contralateral to the neural recordings for state determination. Carprofen (5 mg/kg SC; Putney, Portland, ME) was administered as an anti-inflammatory analgesic.

The skin above the skull was carefully removed and an analgesic (bupivicane; Pfizer, New York, NY) was applied topically to the skull. The skull was then dried with bleach. For P20 rats, any bleeding around the skull was cauterized to ensure that it was completely dry. Vetbond (3M, Minneapolis, MN) was then applied to the skin surrounding the incision, and a custom-manufactured head-plate (Neurotar, Helsinki, Finland) was secured to the skull using cyanoacrylate adhesive.

A trephine drill (1.8 mm; Fine Science Tools, Foster City, CA) was used to drill a hole into the skull above the forelimb representation of M1 (0.5 mm anterior to bregma, 2.2–2.5 mm lateral to the sagittal suture) and either VL (2.0–2.2 mm caudal to bregma, 2.2 mm lateral to the sagittal suture) or VP (2.5–2.8 mm caudal to bregma, 2.5 mm lateral to the sagittal suture). For cerebellar injections, a third hole was drilled above the DCN (3.5 mm caudal to lambda, 2–2.2 mm lateral, contralateral to the other two holes). Care was taken to not puncture the underlying dura matter. A small amount of peanut oil was applied to the dura to prevent desiccation of the underlying tissue. This surgical procedure lasted approximately 30 min.

While recovering from anesthesia, the pup was secured to a custom-made head-fixation block secured to a Mobile HomeCage (NTR000289-01; Neurotar). The height of the pup’s head off the floor was adjusted depending on its age so that, when the pup was exhibiting atonia during active sleep, its body rested on its elbows (P12: 35 mm; P16: 38 mm; P20: 40 mm). EMG wires were carefully secured to ensure they were out of reach of the animal. Pups recovered from anesthesia within 15 minutes and were acclimated for at least 1 h before recording began. By the start of recording, all pups were exhibiting typical behavioral and electrophysiological features of sleep and/or wake (i.e., twitches, grooming, locomotion).

#### Recording Environment

Electrophysiological recordings were performed in a Faraday cage illuminated by flicker-free red LEDs (630 nm; Waveform Lighting, Vancouver, WA). Continuous white noise (70 dB) was present throughout the recording session. To maximize REM sleep, the room’s temperature was maintained between 26.5 and 29° C (Szymusiak and Satinoff, 1981). The animal’s head was positioned away from the entry to the room so that it could not see the experimenter entering or leaving, which was infrequent; whenever possible, the experimenter was outside the room and monitoring the animal and data collection remotely.

#### Electrophysiological Recordings

The nuchal and biceps EMG electrodes were connected to the analogue inputs of a Lab Rat LR-10 acquisition system (Tucker Davis Technologies, Gainesville, FL). The EMG signals were sampled at approximately 1.5 kHz and high-pass filtered at 300 Hz.

Either two 16-channel silicon depth electrodes (A1×16-3mm-100-177-A16 or A1×16-10mm-100-177-A16, depending on the depth of the recording site) or one 32 channel (A2×16-10mm-50-500-177-A16) were coated in fluorescent Dil (Vybrant Dil Cell-Labeling Solution; Life Technologies, Grand Island, NY) before insertion. Electrodes were inserted using a multiprobe manipulator (New Scale Technologies; Victor, NY) controlled by an Xbox controller (Microsoft, Redmond, WA). For M1 recordings, the electrode was inserted tangentially to the cortical surface until all electrode sites were beneath the dura (approximately 1500 μm). For thalamic recordings, the electrode was inserted to a depth ranging from 4500 to 5800 μm, until neurons from VL or VM—which were much more active than neurons in surrounding areas—were centered on the electrode. A chlorinated Ag/Ag-Cl wire (0.25 mm diameter; Medwire, Mt. Vernon, NY) inserted into occipital cortex contralateral to the cortical and thalamic recording sites, served as both reference and ground. Neural signals were sampled at approximately 25 kHz, and both a high-pass filter (0.1 Hz) and a harmonic notch filter (60, 120, and 180 Hz) was applied.

Electrophysiological data were acquired continuously for 3-6 h using SynapseLite (Tucker Davis Technologies). The Mobile HomeCage enabled stable head-fixation while rats were able to locomote, groom, and sleep throughout the recording session.

#### DCN Injections

Pups were prepared for testing as described above. In addition, a microsyringe (1 μL; Hamilton, Reno, NV) was inserted into the DCN contralateral to VL recordings and ipsilateral to the behaviorally scored limbs and whiskers (see Fig. S4A). The recording session began with a 60-min baseline period followed by a 15-min diffusion period and a 240-min recording period. A total volume of 0.5 μL of fluorophore-conjugated muscimol (1.6 mM; Sigma-Aldrich, St. Louis, MO) or saline was injected at a rate of 0.1 μL/min.

#### Video Collection and Synchronization

As described previously (Dooley et al., 2020), video data were synchronized to electrophysiological recordings so that we could better quantify movements and behavioral state. Rats were surrounded by a clear enclosure within the Mobile HomeCage, enabling unimpeded visual access. Video was recorded using a single Blackfly-S camera (FLIR Integrated Systems; Wilsonville, Oregon) positioned at a 45° angle (relative to the pup’s head) and centered on the right forelimb, contralateral to cortical and thalamic recordings. This angle provided an unobstructed view of the right whiskers, forelimb, hindlimb, and tail. Video was collected in SpinView (FLIR Integrated Systems) at 100 frames/s, with a 7 ms exposure time and 720 × 540 pixel resolution.

Video frames were synchronized to the electrophysiological record using an external time-locking stimulus. A red LED, controlled by SynapseLite (Tucker Davis Technologies), was in view of the camera and pulsed every 3 s for a duration of 100 ms. Custom MATLAB scripts determined the number of frames between each LED pulse to check for dropped frames. Although infrequent, when the number of frames between pulses was not equal to 300 (3 s inter-pulse interval × 100 frames/s), a “dummy frame” was inserted in that location. This ensured that the video and electrophysiological data were synchronized to within 10 ms throughout the recording session.

#### Histology

At the end of the recording session, the pup was euthanized with ketamine/xylazine (10:1; >0.08 mg/kg) and perfused with 0.1 M phosphate-buffered saline (PBS) followed by 4% paraformaldehyde (PFA). The brain was extracted and post-fixed in 4% PFA for at least 24 h and was then transferred to a 30% sucrose solution at least 24 h before sectioning.

To confirm the electrode’s location in M1, the brain was either flattened and sectioned tangentially to the pial surface or sectioned coronally. When sectioned tangentially, the right cortical hemisphere was dissected from the subcortical tissue and flattened between two glass slides (separated using a 1.5 mm spacer) for 5-15 min. Small weights (10 g) applied light pressure to the upper slide. To visualize thalamic and cerebellar tissue, brains were always sectioned coronally. Regardless of the plane of section, the tissue was sectioned at 80 μm. Wet-mounted sections were imaged at 2.5× using a fluorescent microscope and digital camera (Leica Microsystems, Buffalo Grove, IL) to identify the location of the DiI.

Cortical and thalamic sections were stained for cytochrome oxidase (CO), which reliably delineates primary sensory areas in cortex and nuclei in thalamus, including VP and VL, at these ages (Cox et al., 1996; Seelke et al., 2012). Briefly, cytochrome C (3 mg per 10 mL solution; Sigma-Aldrich), catalase (2 mg per 10 mL solution; Sigma-Aldrich) and 3,3’-diaminobenzidine tetrahydrochloride (DAB; 5 mg per 10 mL solution; Spectrum, Henderson, NV) were dissolved in a 1:1 dilution of PB-H2O and distilled water. Sections were developed in well plates on a shaker at 35–40°C at approximately 100 rotations per min for 3–6 h, after which they were washed and mounted and allowed at least 48 h to dry. Once dry, sections were placed in citrus clearing solution and cover slipped.

Cerebellar sections were stained for Nissl, which reliably shows the boundaries of the interposed, dentate, and fastigial nuclei by P12 (Del Rio-Bermudez *et al*., 2016). Briefly, cerebellar sections were mounted directly to gelatin-dipped slides and allowed to dry for at least 48 h. Next, sections were rehydrated in distilled H_2_O and soaked in cresyl violet for 45 min. They were then rinsed in deionized H_2_O and transferred to a solution composed of 75% ethanol, 24.5% deionized H2O, and 0.5% acetic acid. Sections were checked several times per minute until they had sufficiently bleached, and then rinsed for three min in progressively higher concentrations of ethanol, placed in citrus clearing solution, and cover slipped.

Stained sections were photographed at 2.5× or 5× magnification. Multiple photographs were combined into a single composite image (Microsoft Image Composite Editor; Microsoft, Redmond, WA) and the electrode or needle location, as well as drug diffusion, were visualized in relation to areal, nuclear, and laminar boundaries of the stained tissue.

#### Classification of Behavioral State

As described previously, behavior, nuchal EMG signals, and cortical LFP were used to identify periods of wake, active sleep, and quiet sleep (Seelke and Blumberg, 2008). Wake was characterized by periods of high nuchal muscle tone and wake-related behaviors (e.g., locomotion, grooming). Active sleep was characterized by the occurrence of myoclonic twitches against a background of muscle atonia. At P16 and P20, active sleep was also accompanied by continuous theta oscillations in thalamus (Gervasoni et al., 2004). During quiet sleep, we observed high cortical delta power, behavioral quiescence, and moderate nuchal muscle tone, although periods of high delta power were sometime accompanied by nuchal muscle atonia. Periods not assigned as active or quiet sleep were designated as wake. Active wake was defined as that part of the wake period that was within 3 s of a movement.

#### Classification of Movement

As described previously (Dooley *et al*., 2020), to quantify periods of movement, we used custom MATLAB scripts to detect frame-by-frame changes in pixel intensity within regions-of-interest (ROIs). The number of pixels within the ROI that exhibited an intensity change >5% were summed. This calculation was performed for each frame, resulting in a time series of quantified movement.

For twitches, the movement timeseries for an ROI containing the body part of interest (e.g., forelimb) was visualized alongside the video using Spike2 (Version 8; Cambridge Electronic Design, Cambridge, UK). Peaks in the timeseries were confirmed to be twitches of the appropriate body part. Twitch onset was defined as the first frame in which movement occurred (as determined by a change in intensity in the ROI). For the forelimb, hindlimb, and tail, this method effectively counted every visible twitch, since even twitches in rapid succession have distinct onset and offset times. However, for whisker twitches, alternating protractions and retractions did not always have clear temporal boundaries, thus making it difficult to determine when one whisker twitch ended and another began. In these instances, only the first whisker twitch in a series of twitches was counted.

For wake movements, an ROI of the entire animal was used. Because multiple wake movements typically occur in quick succession, it was often the case that only the first wake movement in a bout could be used. Thus, our quantification of wake movements underestimates the total number of movements produced.

#### Local Field Potential

For all local field potential analyses, the raw neural signal was down sampled to ~1000 Hz and smoothed using a moving Gaussian kernel with a half-width of 0.5 ms.

#### Spike Sorting

SynapseLite files were converted to binary files using custom Matlab scripts and sorted with Kilosort2 (Pachitariu et al., 2016). Briefly, data were whitened (covariance-standardized) and band-pass filtered (300-5000 Hz) before spike detection. Template-matching was implemented to sort the event waveforms into clusters. The first-pass spike detection threshold was set to six standard deviations below the mean and the second-pass threshold was set to five standard deviations below the mean. The minimum allowable firing rate was set to 0.01 spikes/s and the bin size for template detection was set to 262,400 sample points, or approximately 11 s. All other Kilosort2 parameters were left at default values.

Clusters were visualized and sorted in Phy2 (Rossant and Harris, 2013). Putative single units had spike waveforms that reliably fit within a well-isolated waveform template, appeared in a principal component analysis as a distinct cluster, and had an auto-correlogram with a decreased firing rate at a time lag of 0 (indicative of a unit’s refractory period).

Clusters meeting the first two criteria but not the third were considered multi-units and were discarded from analysis. Any putative unit with a waveform template indicative of electrical noise, a firing rate < 0.01 spikes/s, or amplitude drift across the recording period was discarded.

### QUANTIFICATION AND STATISTICAL ANALYSIS

#### Determination of Twitch-Responsiveness

All analyses of neural data were performed in MATLAB using custom-written scripts. The relation between neural activity and twitches was assessed as follows. First, all twitches of each body part (forelimb, hindlimb, whiskers, tail) were behaviorally scored. For each body part that had more than 20 twitches, perievent histograms of neural activity were constructed (window size: −3 to 3 s; bin size: 10 ms). Next, we determined the mean baseline firing rate from −3 to −0.5 s. Finally, we z-scored the perievent histograms by subtracting the baseline from the raw perievent histograms and dividing this value by the standard deviation of the baseline. A neuron was considered “responsive” to twitches if it showed a clear peak with a z-score of at least 3.5 for at least one body part.

To accurately determine a neuron’s preferred somatotopic response (see Figs. 2A, S2C), we only analyzed twitches for a given body part that were separated by at least 100 ms from twitches of other body parts. The preferred body part for each neuron was the body part with the largest peak in the z-scored perievent histogram (see Figs. 2B, S4E).

#### Movement Analysis

For each movement type (twitches of different body parts and wake movements), we calculated the average displacement produced by that movement for each pup (see Fig. S2D). To ensure subsequent twitches (of the same or different body parts) did not influence the average displacement of a twitch, for this analysis, we used the same subset of twitch triggers used in the somatotopy analysis above (i.e., twitches with inter-twitch-intervals of at least 100 ms). Median average displacement was calculated over all triggered twitches (or wake movements) at each timepoint. Because the size of the ROI used to assess movements varied across age and body part, median values for each animal were always normalized.

#### Width at Half-Height

To measure width at half-height, the displacement and neural data were first smoothed using a 5-bin kernel and then upscaled from 10 ms bins to 1 ms bins using the interp1() function in MATLAB. The data were then normalized so that the baseline was equal to 0 and the peak was set to 1. The width at half-height (in ms) equaled the number of bins between 0.5 and 1.

#### Statistical Analyses

All statistical tests were performed using MATLAB. Alpha was set at 0.05 for all analyses; when appropriate, the Bonferroni procedure was used to correct for multiple comparisons. Unless otherwise stated, mean data are always reported with their standard error (SE). Normally distributed data were tested for significance using a one-way ANOVA, two-way ANOVA, or t test. Categorical data were tested for significance using a Chi-squared test.

Non-normally distributed data were visualized using violin plots, with the red line signifying the median. Two-group data were tested for significance using the Kruskal-Wallis nonparametric test. For individual groups, a Wilcoxon signed-rank test was used. Violin plots were constructed absent outliers, which were determined using the isoutlier() function in MATLAB. This function excludes datapoints that are more than three scaled median-absolute-deviations from the median.

---

**Figure S1.**
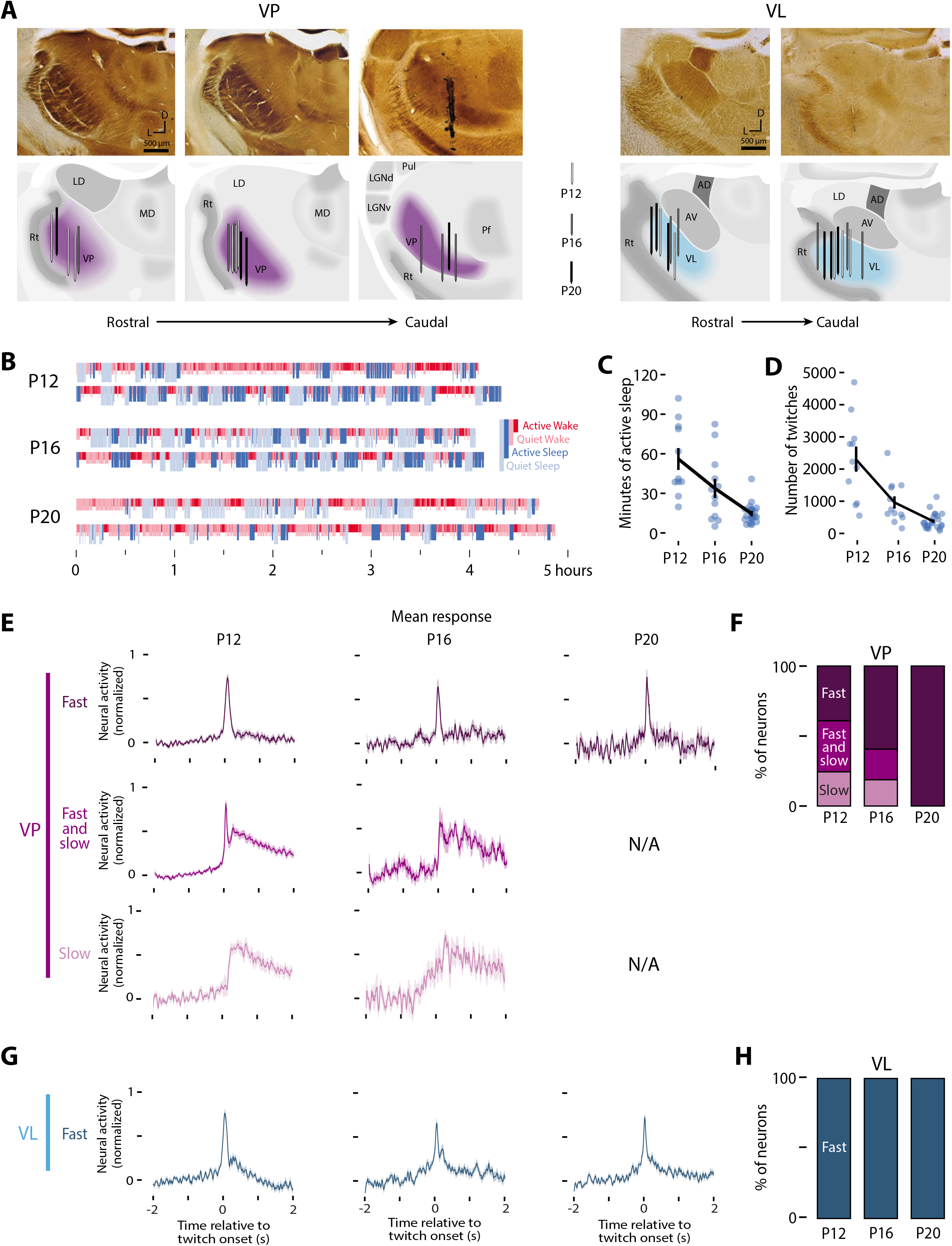
Anatomical and behavioral details, Related to Figure 1. (A) *Top*: Representative tissue stained for CO showing the rostral to caudal progression of VP (left) and VL (right). VP was differentiated from the surrounding nuclei by its dark appearance in CO-stained tissue. VL was located rostral to VP and appeared lighter than adjacent nuclei (Rt, AV, and VP) in CO-stained tissue. *Bottom*: Location of the recording electrodes in VP (left, purple) and VL (right, cyan) in rats at P12 (white electrodes), P16 (gray electrodes), and P20 (black electrodes). CO: Cytochrome oxidase. LD: Lateral dorsal nucleus. MD: Medial dorsal nucleus. Rt: Thalamic reticular nucleus. VP: Ventral posterior nucleus. Pul: Visual pulvinar. LGNd/v: Lateral geniculate nucleus, dorsal and ventral divisions. Pf: Parafascicular nucleus. AD: Anterodorsal nucleus. AV: Anteroventral nucleus. VL: Ventrolateral nucleus. (B) Representative sleep staging throughout the recording session for two rats at each age. Periods of active wake (dark red), quiet wake (light red), active sleep (dark blue), and quiet sleep (light blue) are shown. (C) Mean (± SE) number of minutes of active sleep for each animal across each recording session. Data for individual pups are also shown. The amount of time spent in active sleep decreased significantly across age (F2,44 = 18.6, p < 0.0001). (D) Mean (± SE) number of twitches for each animal across each recording session. Data for individual pups are also shown. Twitching decreased significantly across age (F2,44 = 26.8, p < 0.0001). (E) Mean (± SE) normalized neural response of VP neurons categorized as fast (top, dark purple), fast and slow (middle, purple), and slow (bottom, light purple) at P12, P16, and P20. N/A indicates an absence of neurons in that category. (F) Stacked plots showing the percentage of VP neurons categorized as fast, fast and slow, and slow at each age. (G) Same as in (E), but for VL, for which only fast neurons were observed (dark cyan). (H) Same as in (F), but for VL.

**Figure S2.**
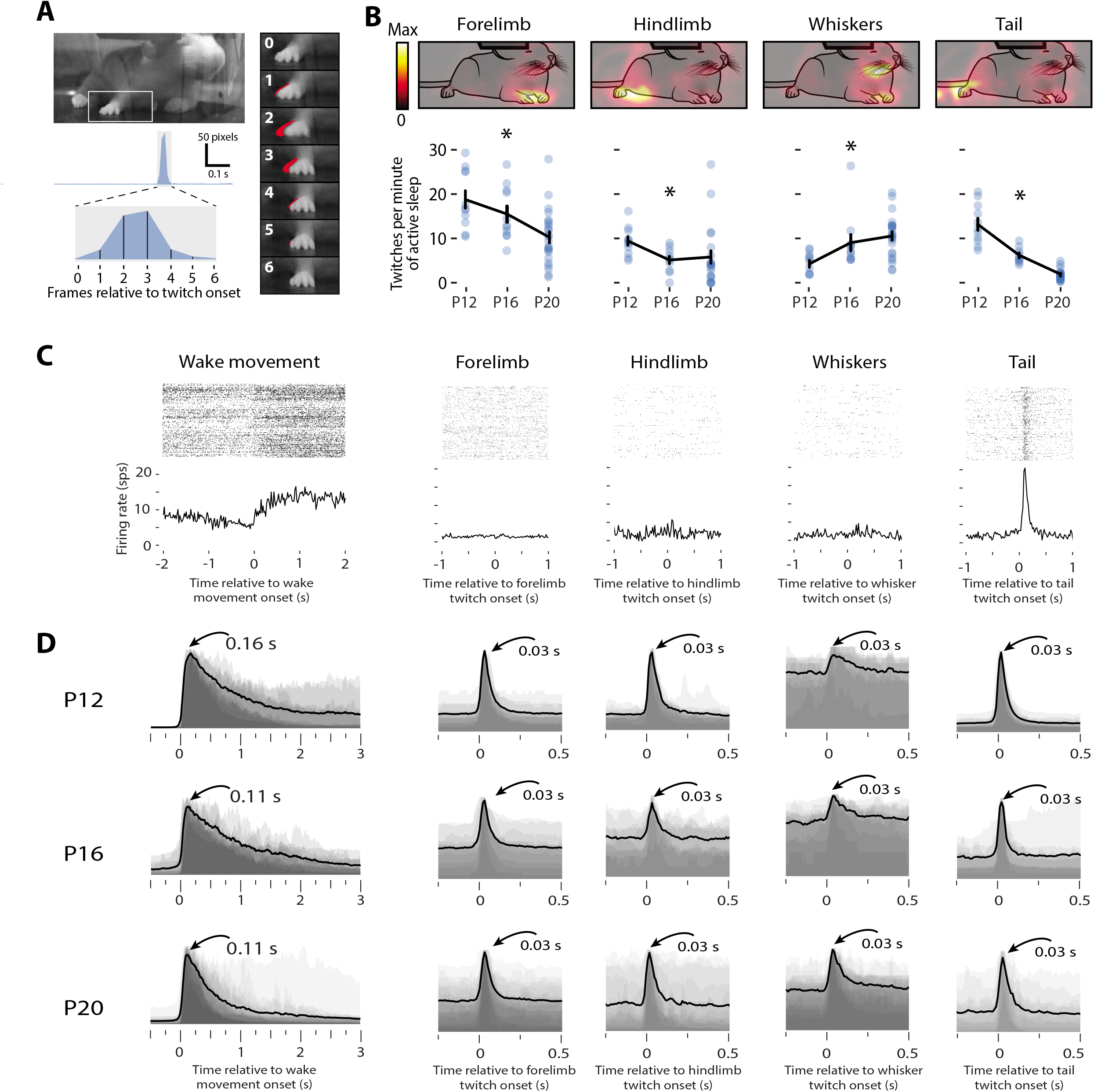
Twitch rates, somatotopy, and kinematics across development, Related to Figure 2. (A) The image at top-left shows a single frame of a P12 rat just before the onset of a twitch of the forelimb. The area within the white rectangle is shown across seven frames in the panels at right. Starting with the top-most frame (frame 0) and moving down frame by frame, the pixels that change from the previous frame are highlighted in red. The forelimb moves the most in frames 2 and 3 and, by frame 6, the movement has stopped. At left-middle, the output of the twitch-movement analysis for a 1-s interval of active sleep. The region around the twitch (gray box) is expanded below to show the time-course over the seven frames. (B) *Top*: Drawing of a P20 rat overlaid with a heatmap showing the pixels where movement was most likely to occur in the ten frames after the onset of forelimb, hindlimb, whiskers, and tail twitches. *Bottom*: Mean (± SE) rate of twitches per min of active sleep for twitches of the forelimb, hindlimb, whiskers, and tail across age. Data for individual sessions are also shown. The rate of twitching decreased significantly across age for the forelimbs (F_2,44_ = 8.01, p < 0.005), hindlimbs (F_2,39_ = 7.61, p < 0.005), and tail (F_2,44_ = 62.9, p < 0.0001), but increased for the whiskers (F_2,43_ = 5.46, p < 0.01). (C) Perievent histograms for a representative VP neuron at P12 for movements during wake (single panel at left) and twitches during sleep (four panels at right). This neuron responds weakly to wake movements and negligibly to twitches of the forelimb, hindlimb, and whiskers. In contrast, this neuron responses strongly to twitches of the tail, reflecting somatotopic precision. (D) Mean displacement produced during wake movements, as well as twitches of the forelimb, hindlimb, whiskers, and tail across age. Each shaded gray region denotes the normalized median displacement produced by that movement for an individual pup. The black line is the mean across all pups for that body part and age. For all twitches across all ages, the mean peak displacement occurs 0.03 s (or three frames) after twitch onset.

**Figure S3.**
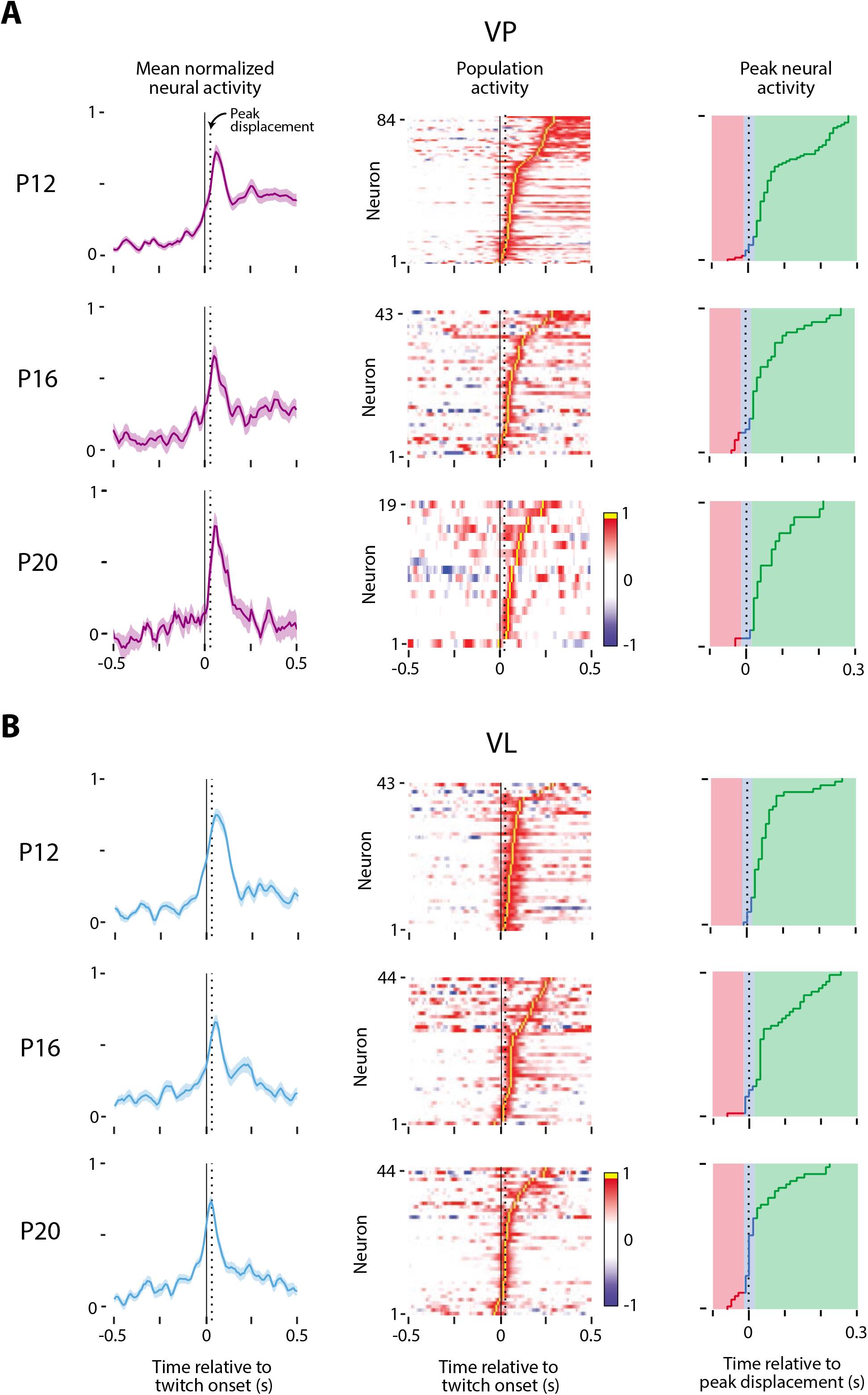
Population-level twitch-related activity in VP and VL, Related to Figure 3. (A) *Left column*: Mean (± SE) normalized neural activity of twitch-responsive VP neurons across age relative to twitch onset (solid vertical line). Peak twitch displacement, occurring at 0.03 s after twitch onset, is also shown (dotted vertical line). *Middle column*: Heatmap of each twitch-responsive VP neuron’s activity relative to a twitch, sorted from bottom by increasing peak time. Each neuron’s peak activity (yellow) is represented in the right column. *Right column*: The timing of peak activity for each neuron plotted on top of the categories used to classify neural responses in Fig. 3A. Red indicates a neuron that fires before peak displacement, whereas green indicates a neuron that fires after peak displacement. Blue are the neurons that fire within 10 ms of peak displacement. (B) Same as in (A), but for VL.

**Figure S4.**
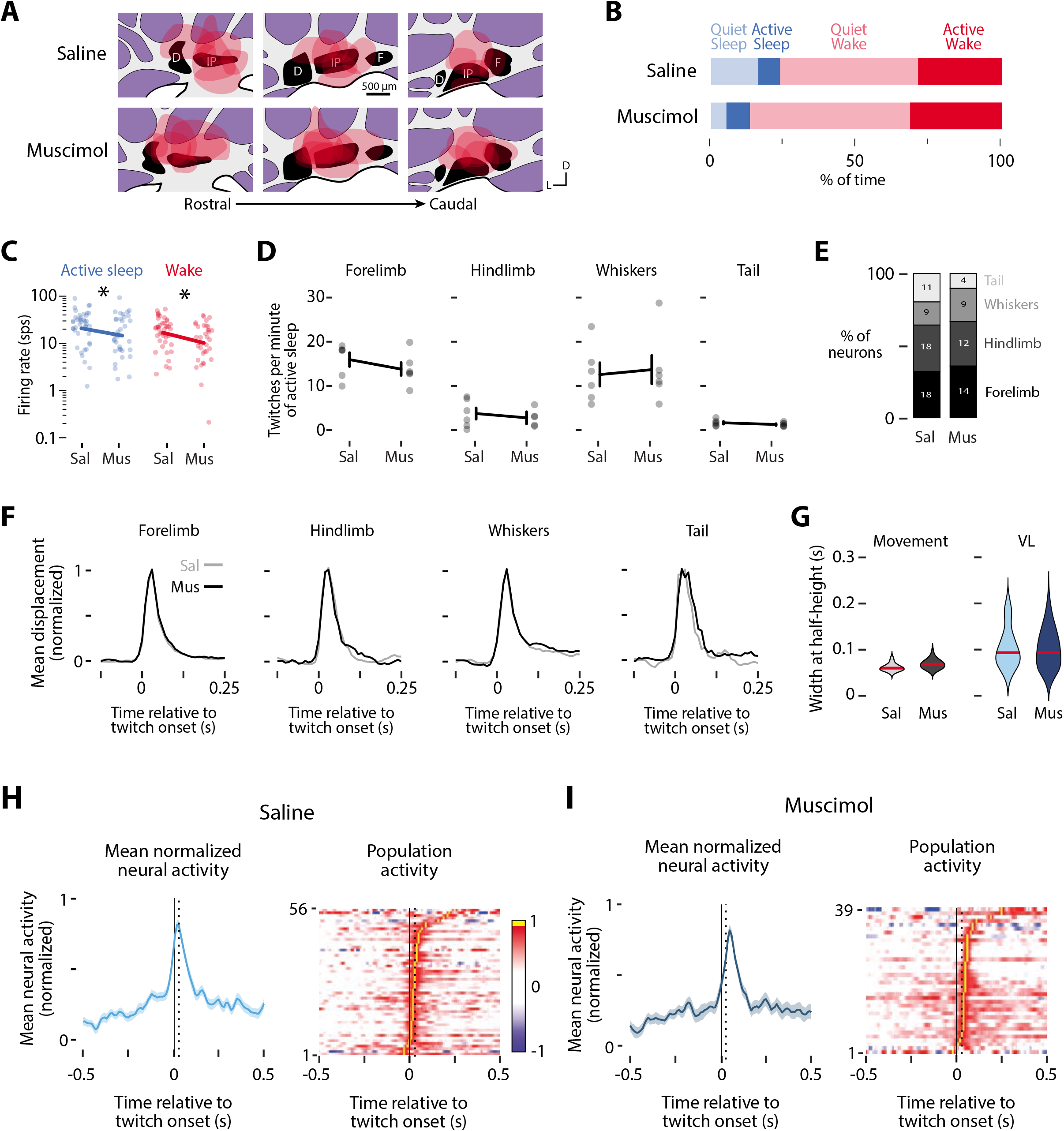
Twitch-related VL activity after injection of muscimol or saline into the DCN at P20, Related to Figure 4. (A) Drawings of the DCN nuclei across three coronal sections to show the lateral and rostrocaudal diffusion of saline (top) or muscimol (bottom). N=6 per group. D: Dentate. IP: Interposed. F: Fastigial. (B) Mean percentage of time spent in quiet sleep (light blue), active sleep (dark blue), quiet wake (light red), and active wake (dark red) for pups injected with saline or muscimol. (C) Mean log-normalized firing rate of VL neurons during active sleep (blue, left) and wake (red, right) following injection of saline (Sal) or muscimol (Mus). Data for individual neurons are also shown. A 2-way ANOVA reveals a significant effect of injection type (F_1,152_ = 8.42, p < 0.005) but not behavioral state, with muscimol-injected pups showing less neural activity compared to saline animals (denoted by asterisk). (D) Mean (± SEM) rate of twitches per min of active sleep for twitches of the forelimb, hindlimb, whiskers, and tail across age. Data for individual pups are also shown. None of the differences are statistically significant. (E) Stacked plots showing the percentage of twitch-responsive VL neurons that respond to twitches of different body part in pups injected with saline (left) or muscimol (right). The number of neurons is also indicated. (F) Mean normalized movement displacement during twitches of the forelimb, hindlimb, whiskers, and tail in pups injected with saline (gray lines) or muscimol (black lines). The displacement profiles across groups are nearly identical. (G) *Left*: Mean (± SE) normalized VL activity for saline-injected pups. Peak displacement (0.03 s after twitch onset) is shown as a dotted line. *Right*: Heatmap showing each twitch-responsive neuron’s activity relative to a twitch, sorted by peak time. Each neuron’s peak activity (yellow) is represented in Fig. 4B. The timing of this peak activity relative to peak displacement (dotted line) is used to classify neural responses in Fig. 4C. (H) Same as in (G), but for muscimol-injected pups. (I) *Left*: Violin plots of the width at half-height for twitch movements for pups injected with saline or muscimol. *Right*: Violin plots of the width at half-height for twitch-responsive VL neurons for pups injected with saline or muscimol. The solid red lines denote median values. Neither of the group differences is significant.

## Notes

### Competing Interest Statement

The authors have declared no competing interest.

